# Topography modulates climate sensitivity of multidecadal trends of holm oak decline

**DOI:** 10.1101/2023.03.09.531879

**Authors:** Ana López-Ballesteros, Emilio Rodríguez-Caballero, Gerardo Moreno, Paula Escribano, Ana María Hereş, Jorge Curiel-Yuste

**Author notes:** Corresponding author: Ana López-Ballesteros.

## Abstract

Forest decline events have increased worldwide over the last decades, being holm oak one of the tree species with the most worrying trends across Europe. Previous research identified drought and soil pathogens as the main causes behind holm oak decline. However, despite tree health loss is a multifactorial phenomenon where abiotic and biotic factors interact in time and space, there are some abiotic factors whose influence has been commonly overlooked. Here, we evaluate how land use (forests versus savannas), topography, and climate extremes jointly relate to spatiotemporal patterns of holm oak defoliation over almost three decades (1987-2014) in Spain, where holm oak represents the 25% of the national forested area. We found an increasing defoliation trend in 119 of the total 134 holm oak plots evaluated, being this increase rate significantly higher in forests compared to savannas. Moreover, we have detected that the interaction between topography and summer drought can explain trends of holm oak decline across the Mediterranean region. While a higher occurrence of dry summers increases defoliation trends in complex terrains where forests dominate, an inverse relationship was found at flatter terrains where savannas are located. Our results contribute to growing evidence of the influence of local topography, tightly linked to potential soil water availability, on forest functioning, as it can shape forest vulnerability against climate extremes. The present work could assist the identification of potential tree decline hotspots over the Mediterranean region. Moreover, our findings suggest that forest adaptive management will be key to guarantee the health and future stability of Mediterranean oak ecosystems, especially in the topographically more complex areas where tree vulnerability to climate extremes may be greater.

## 1. Introduction

The number of tree mortality and forest decline events have increased worldwide over the last decades (Allen et al., 2015; Hammond et al., 2022), with accelerating rates being associated to climate change (McDowell et al., 2022; Parmesan et al., 2022). These decline pattens have global implications in the current climate change context as they negatively affect the global land carbon sink (Clark et al., 2010; Haberstroh et al., 2022; Liu et al., 2023). Although the primary physiological mechanisms behind tree mortality are related to water and heat stress (Allen et al., 2010, 2015), climate extremes can also amplify the occurrence and impact of other disturbance drivers, such as wildfires and pathogen outbreaks (Brando et al., 2014; Gea-Izquierdo et al., 2021; Seidl et al., 2017). Regional studies based on either systematic forest inventories or remote sensing observations have identified the Mediterranean basin as a hotspot of forest decline (Carnicer et al., 2011; Forest, 2020; Senf et al., 2020). This has coincided with an increase in the magnitude and frequency of heat waves and droughts since the pre-Industrial era (Douville et al., 2022; Markonis et al., 2021; Vicente-Serrano et al., 2014; Seneviratne et al., 2022). Foreseeably, the higher occurrence probability of longer and more severe droughts projected across the Mediterranean region (García-Valdecasas Ojeda et al., 2021) will pose an even higher mortality risk to Mediterranean tree species in the near future.

One of these tree species that have already shown clear signs of decline is Quercus ilex L. (hereinafter holm oak). Holm oak naturally grows in areas with Mediterranean climate characterized by summer-drought periods and cold or mild moist winters (Lionello et al., 2006). This species occurs over a large range of altitudes, from sea-level up to 2500 m.a.s.l. (Barbero et al., 1992), given its ability to cope with drought and heat stress by means of stomatal regulation and photoprotective mechanisms (Alonso-Forn et al., 2021). However, acute signs of holm oak decline have been detected across de Iberian Peninsula, including growth reduction (Corcuera et al., 2004; Gea-Izquierdo et al., 2011; Hereş et al., 2018), partial crown defoliation (Carnicer et al., 2011; FOREST, 2020), branch dieback, and tree death (Lloret et al., 2004; Sánchez-Cuesta et al., 2021). In fact, Europe-wide defoliation surveys have identified evergreen oaks as the species group with the highest increase in defoliation over the last 20 years (Michel et al., 2021). These declining symptoms have been detected in both major land uses where holm oak dominates across southern Europe: coppice forests and wood pastures, which differ in both stand structure and management (Gazol et al., 2020). Coppices refer to Mediterranean holm oak forests with high tree density and multi-stemmed structure that have traditionally been managed to provide firewood and charcoal given the high resprouting capability of this tree species after cutting (Giovannini et al., 1992). Instead, holm-oak wood pastures – known as dehesas or montados in Spain and Portugal, respectively – are human-made savanna-like ecosystems that have been conventionally managed as agroforestry systems where the low tree density allows for extensive livestock farming (Moreno & Rolo, 2019; Plieninger et al., 2021; Pulido et al., 2001). The geographical distribution of these landuse types is generally determined by local topography. While wood pastures usually occupy topographically smooth areas, coppice forests are largely found in topographically complex areas with steeper slopes and shallower and drier soils. Understanding the causes of holm oak decline at both forest- and savanna-like ecosystems remains a scientific priority to preserve the valuable socio-ecological services that these habitats have provided for centuries (Gil-Pelegrín et al., 2017; Plieninger et al., 2015; Sáez et al., 2007; Stavi et al., 2022; Terradas, 1999).

Previous research suggests different prevailing mechanisms behind holm oak decline in these land-use types. On one hand, decline episodes in holm oak forests have been associated to carbon starvation (Galiano et al., 2012), and drought-induced xylem embolism, which reduces the water uptake capacity of the trees leading to leaf desiccation, and shoot, branch, or tree mortality (Barbeta et al., 2013; Corcuera et al., 2004; Lloret et al., 2004). Furthermore, the abandonment of the traditional coppicing practices derived from rural depopulation (B. C. López et al., 2009; Serrada et al., 1992) has shaped the structure of these forests into stands with a higher density of overaged individuals with aggravated water stress and drought vulnerability (Rodríguez-Calcerrada et al., 2011, Gazol et al., 2020). On the other hand, holm oak decline in Mediterranean savannas has been primarily associated to soil-borne pathogen outbreaks of Phytophtora cinamomii Rands (Brasier et al., 1993), which causes hydraulic failure through root damage, and triggers similar decline symptoms to the drought-induced ones (Corcobado et al., 2014; de Sampaio e Paiva Camilo-Alves et al., 2013; Encinas-Valero et al., 2021). Recent evidence indicates that defoliation patterns in savannas may also be caused by changes in shot-to-root non-structural carbon allocation patterns associated with the belowground stress caused by the pathogen (Encinas-Valero et al., 2022). However, there is no consensus on how pathogenicity and climate stress interact. While some studies state that pathogenicity can be presumably amplified by climatic stress (Brasier, 1996; González et al., 2020; Rodríguez-Molina et al., 2005), others suggest that drought reduces the risk of pathogen damage (Homet et al., 2019; Serrano et al., 2022).

These contrasting results reveals that holm oak health loss and mortality is a multifactorial phenomenon where abiotic and biotic factors interact in time and space (Brando et al., 2014; Gea-Izquierdo et al., 2021; Seidl et al., 2017), but not all factors potentially involved have been equally evaluated. It has been demonstrated that land use (i.e. forest structure and management) can modulate the effect of climate on forest die-back/die-off/mortality (Alfaro-Sánchez et al., 2019; Gentilesca et al., 2017; Sangüesa-Barreda et al., 2015). Still, very few studies have assessed the role of this factor in holm oak decline (e.g., Gazol et al., 2020). Likewise, previous research has shown that local topography influences how forest resilience (Carnicer et al., 2021), phenology (Adams et al., 2021), and decline (Wang et al., 2021) respond to climate variability. In this sense, topographic features together with vegetation structure strongly determine local microclimate (Bennie et al., 2008; Bramer et al., 2018; Ma et al., 2010; Scherrer & Körner, 2010) and soil water availability (Burt & Butcher, 1985; Huang et al., 2012; Tromp-van Meerveld & McDonnell, 2006). Despite the complex topography of the Mediterranean region, there is a lack of regional studies where systematic observations of holm oak decline cover areas with topographic variability and where both holm oak forests and savannas are represented.

In this study, our general aim is to better understand the phenomenon of holm oak decline, given its socio-ecological and economic relevance in the Mediterranean basin and especially in Spain, where holm oak represents the 25% of the forested area and the 16% of wood production (Terradas, 1999). Thus, we have made use of public databases to evaluate how land use, topography, and climate extremes jointly relate to spatiotemporal patterns of holm oak defoliation over almost three decades (1987-2014) in Spain. We hypothesize that both, land use and associated topographic features determine the vulnerability of holm oak to climate variability and extremes, resulting in different climate-decline relationships in holm oak-dominated ecosystems. Our specific objectives are: (i) to compare multidecadal trends of holm oak defoliation between the main land-use types (i.e., forests vs savannas); and (ii) to disentangle the interactive influence of land use, local topography and climate anomalies on multidecadal trends of holm oak defoliation.

## 2. Material and methods

### 2.1. Data sources and processing

Several public datasets were utilized to investigate the influence of land use, climate, and topography on holm oak defoliation across Spain over the period 1987-2014. Main specifications of these geospatial and survey datasets are summarized in Table S1.

#### 2.1.1. Defoliation data

The annual holm oak crown defoliation data used in this study spans from 1987 to 2014 and was obtained from the survey data of the Level 1 network of the International Cooperative Programme on Assessment and Monitoring of Air Pollution Effects on Forests (ICP Forests; Eichhorn et al. 2016). Starting from 1987, the Level I monitoring scheme of ICP Forests annually surveys crown defoliation at >5000 forested plots, which are systematically arranged in a nominal 16×16km grid throughout Europe. For this study, we only used plots where holm oak was present, and a minimum number of three observations every 7-yr period were available over the whole 28-yr study period (1987-2014). This filtering criteria yielded a total of 134 plots spread across the entire holm oak distribution area in Spain (Fig. 1) with data spanning from 24 to 28 years surveyed. These circular plots of 50-m radius contained at least one predominant, dominant, and co-dominant holm oak tree (i.e., minimum height of 60 cm) that showed no significant mechanical damage (Eichhorn et al. 2016). From the total of 134 plots analyzed in this study, 90% of them included at least five holm oak trees per plot. Tree crown defoliation was visually evaluated, always during the summer season, by associating a percentage of leaf loss in the assessable crown as compared to a reference tree, using a sliding scale of 5%. Reference trees were defined as trees with the best full crown foliage that could grow at each plot.

**Figure 1.**
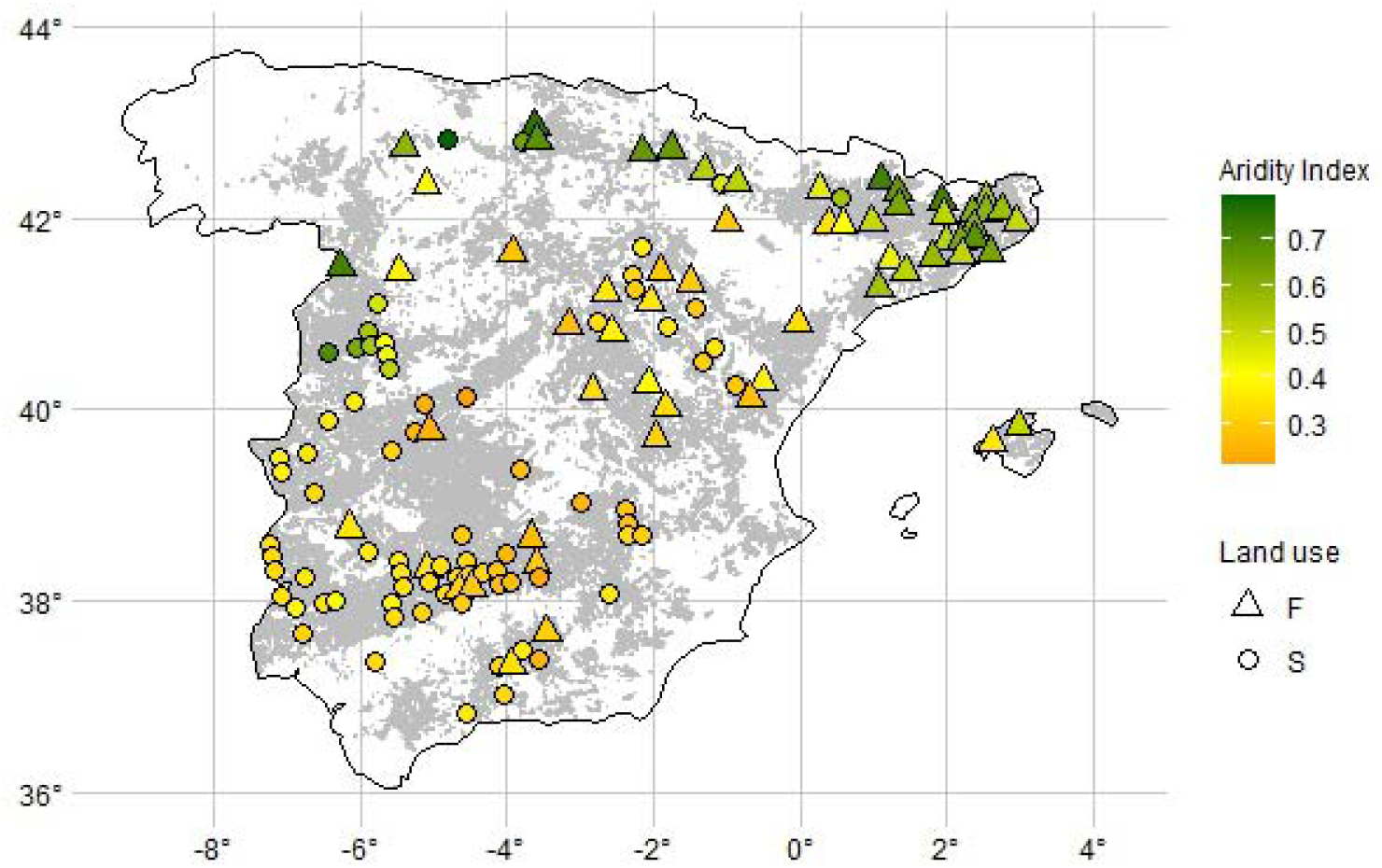
Location of the 134 ICP holm oak plots over the distribution area of holm oak in Spain (grey area). Circles and triangles represent savanna (S) and forest (F) plots, respectively, while their colors indicate the annual averaged plot-mean aridity index (AI) over the 1970-2000 period. This plot-mean AI was extracted from the Global Aridity Index and Potential Evapo-Transpiration (ET°) Climate Database v2 (Trabucco and Zomer, 2019). The holm oak distribution map was provided by GIS-FOREST INIA (Ministerio de Medio Ambiente, y medio Rural y Marino, 2009).

#### 2.1.2. Land use data

The land use of each of the selected ICP plots was derived from the 2014 version (last version available) of the SIOSE (Land Cover and Land Use Information System of Spain) database (1:25000) (Equipo Técnico Nacional SIOSE, 2015a, 2015b). We extracted the codes of those land use polygons that overlap the centroids of each of the 134 ICP holm oak plots. Land use codes provide information of the cover types present in each polygon together with their respective coverage area over the total polygon area. Based on this information, we defined two different land use classes, forest (F) and savanna (S). Concretely, when conifer, deciduous and/or evergreen broadleaf forests occupy at least 50% of the polygon, the overlapping ICP plot was defined as a forest, otherwise, it was defined as a savanna. To do so, we used the sf (Pebesma, 2018) R package, and Google Earth Pro (version 7.3.6.9285) to visually double-check assigned land use classes.

#### 2.1.3. Climatic data

Two different databases were utilized to collate climatic data for the ICP holm oak plots utilized in this study. Firstly, the annual average of the aridity index (AI) over the 1970-2000 period was extracted from the Global Aridity Index and Potential Evapo-Transpiration (ET0) Climate Database v2 (1-km spatial resolution; Trabucco and Zomer, 2019). Seasonal maximum, minimum and mean air temperature together with precipitation values were obtained from the Spain02 v5.0 gridded datasets (10-km spatial resolution; Herrera et al., 2012, 2016). Seasonal climatic anomalies (MAM for spring, JJA for summer, SON for autumn, and DJF for winter) were calculated by subtracting the averaged seasonal value over the whole study period (1987-2014) to the seasonally averaged value of each year. Hence, negative anomalies correspond to seasons when temperature and/or precipitation were lower than the historical average, while positive anomalies correspond to seasons when the opposite occurred, i.e. temperature and/or precipitation were higher than the historical average. Anomaly units were degrees Celsius, and mm for temperature, and precipitation, respectively. As defoliation surveys were annually performed during the summer season, summer and autumn of the years when defoliation was measured were excluded from the correlation analysis. Thus, five seasons were analyzed from spring of the previous year to spring of the current year when defoliation measurements were collected. All these data processing steps were performed by using the lubridate (Grolemund & Wickham, 2011), ncdf4 (Pierce, 2019), raster (Hijmans, 2021), sf (Pebesma, 2018), and tidyverse (Wickman et al. 2019) R packages.

#### 2.1.4. Topographic data

Mean and variance values of several topographic variables were obtained for the ICP holm oak plots to consider both the averaged geomorphology of the surface together with its complexity in the posterior statistical analyses. Slope and altitude values were extracted from a 5-m digital elevation model (DEM) obtained from LIDAR flights 1st Coverage of the National Plan for Aerial orthophotography (PNOA; Spanish Geographic Information National Center, CNIG). Mean aspect was derived from the DEM tiles and reclassified into eight groups by using 45-degrees intervals spanning from north to northwest directions. Terrain curvature parameters and topographic wetness index (TWI) were also derived from the DEM tiles to consider plot-surface water flow dynamics in further statistical analyses. Concretely, profile curvature represents the acceleration of the water flow in the slope direction with positive values associated to upwardly convex surface and negative values to upwardly concave surface for a specific grid cell (Florinsky, 2000). Planform curvature represents the convergence of the water flow in the maximum slope-perpendicular direction with negative and positive values representing sidewardly concave and convex surfaces, respectively, while zero values represent linear surface. The TWI quantifies terrain-driven variation in soil moisture by integrating the water supply from upslope catchment area and the slope gradient for each cell in a DEM (Kopecký et al., 2021). To get plot-scale TWI estimates, a sink removal algorithm with a minimum slope threshold of 0.01 degrees was firstly applied to fill the sinks in the DEM tiles prior to calculation of the upslope accumulated area according to the triangular multiple flow direction algorithm (Seibert & McGlynn, 2007). Then, specific catchment area (SCA; m) was computed via the multiple flow direction algorithm (Quinn et al., 1991) and utilized to get the TWI (dimensionless) according to (Beven & Kirkby, 1979), as follows:

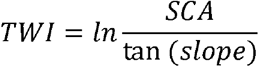

Prior to the extraction of zonal statistics, each raster file was reclassified to assure that the values of each topographic variable were within plausible ranges (see Table S2). All the DEM-derived variables were computed for each tile and subsequently extracted for each ICP plot by using the exactextractr (Baston, 2022), RSAGA (Brenning et al. 2018), raster (Hijmans, 2021), and sf (Pebesma, 2018) R packages.

### 2.2. Statistical analyses

The complete analysis workflow followed in this study is depicted in Fig. 2. Firstly, mean defoliation per plot and year was calculated. Afterwards, multidecadal defoliation trends for each ICP holm oak plot were computed as the Sen’s slope regression coefficient (Sen, 1968) between year and annual defoliation values by using the trend R package (Pohlert, 2020) as this non-parametric slope estimate has been proven more robust for observational data compared to linear regression. Mean and trend values of defoliation over the study period (1987-2014) were then compared between forest and savanna land uses by using either ANOVA or Krustal Wallis tests after checking the normality assumption of the data distribution via a Shapiro normality test and graphic representation. In case of ANOVA test, assumptions of normality of residuals and homogeinity of variance were tested graphically and via Shapiro and Levene’s test, respectively. Eta squared (η2) was computed by means of the rstatix R package (Kassambara, 2022) to provide an estimate of how much variance in the response variable is explained by the explanatory variable.

**Figure 2.**
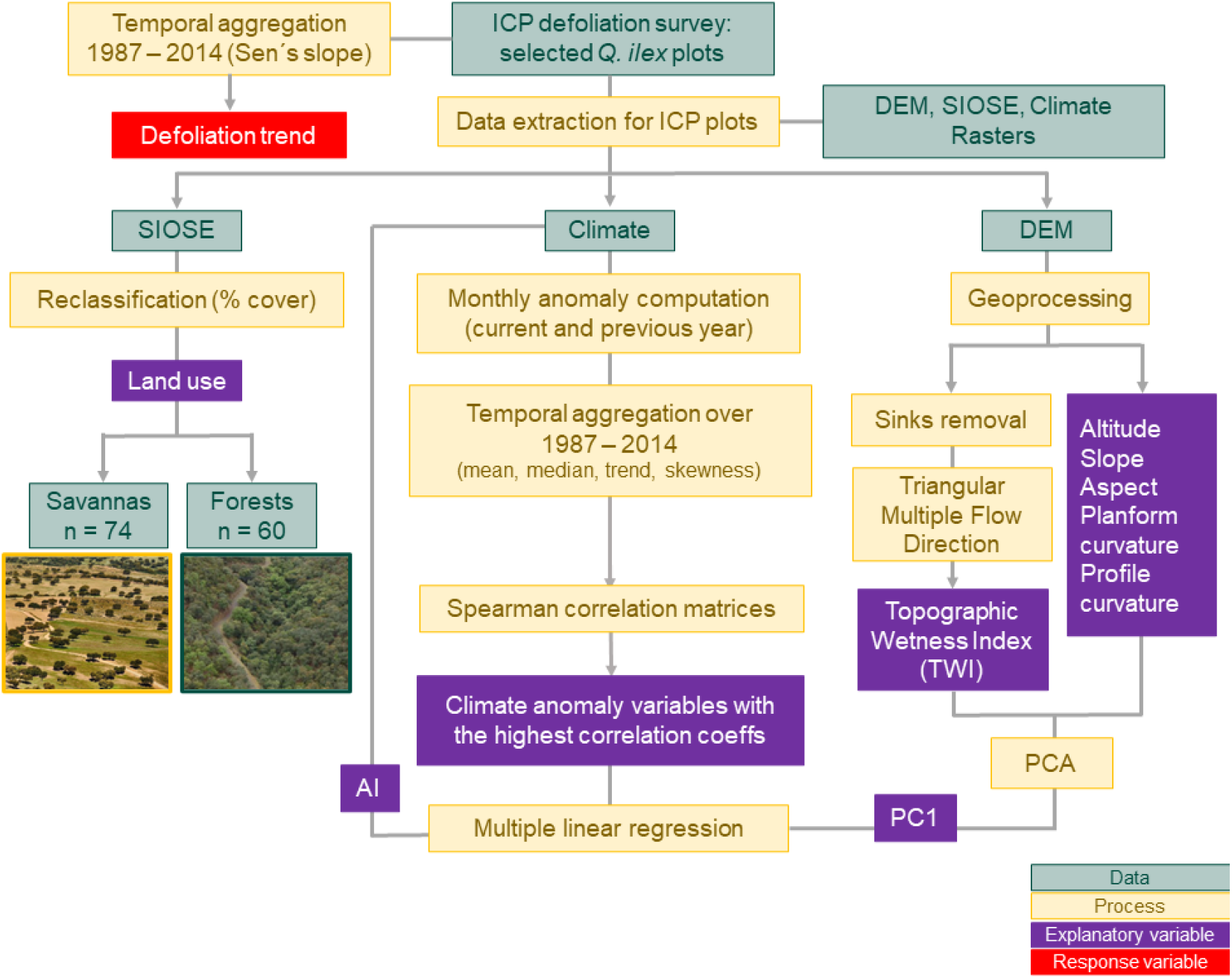
Diagram summarizing the methodological workflow followed in this study.

Multiple linear regression models were used to elucidate causal links between abiotic factors (climate, topography, and land use) and defoliation trends. Prior to modelling, we performed principal component analysis (PCA) to reduce the dimensionality of the topographic characterization data from each plot via the factoextra R package (Kassambara and Mundt, 2020). Variables considered in the PCA corresponded to mean and variance values of altitude, slope, profile and planform curvature, topographic wetness index, and reclassified mean aspect. Moreover, seasonal climate anomalies were temporally aggregated by computing the mean, median, skewness, and Sen’s slope values over the study period (1987-2014). Then, Spearman correlation matrices among aggregated climatic anomalies, and between these and defoliation trends were computed to select those climatic variables to be included in the defoliation model while avoiding multicollinearity. The Spearman coefficients, together with their associated p-values, were calculated by using the Hmisc R package (Harrell, 2021). The defoliation model was built via multiple linear regression. The defoliation Sen’s slopes were set as the response variable while explanatory variables of the full model corresponded to the topographic features synthesized via PCA, the previously selected aggregated seasonal climate anomalies, the interactions among them, and the aridity index (Fig. 2). All these explanatory variables were scaled prior to modelling. Model selection was performed by using the backward and forward stepwise Akaike’s information criterion (AIC) algorithm (Venables & Ripley, 2002). The residuals of the final model were evaluated to check whether the normality and heteroskedasticity assumptions were fulfilled. Spatial autocorrelation was checked by means of the Moran’s I test for distance-based autocorrelation (Moran, 1950). The R packages utilized to perform statistical analyses were car (Fox and Weisberg, 2019), DHARMa (Hartig, 2022), effects (Fox and Weisberg, 2018, 2019), interactions (Long, 2019), jtools (Long, 2022), MASS (Venables & Ripley, 2002), and performance (Lüdecke et al., 2021). The significance level of all the statistical analyses performed in this study was 95% (α = 0.05). For all the analyses explained in material and methods section, data curation and visualization were performed via the tidyverse collection of R packages (Wickman et al. 2019) in R software (version 4.2.2.).

## 3. Results

The obtained results show that crown tree defoliation of holm oak across Spain has increased over the 28-yr study period regardless of the land-use type at an average rate of 0.41 % year-1 (Fig. 3). Specifically, in 119 of the total 134 holm oak plots an increasing defoliation trend was found (i.e. Sen’s slope above 0). Mean annual crown tree defoliation equated to 19.43% with no differences between land uses based on the Krustal Wallis test results (p-value = 0.0669; chi-squared = 3.3572; Fig. 4a). Though, the increase of crown defoliation was more acute in forests (n = 60) than in savannas (n = 74) based on the one-way ANOVA test results (p-value = 0.0012; F value = 10.93; η2 = 0.08; Fig. 4b).

**Figure 3.**
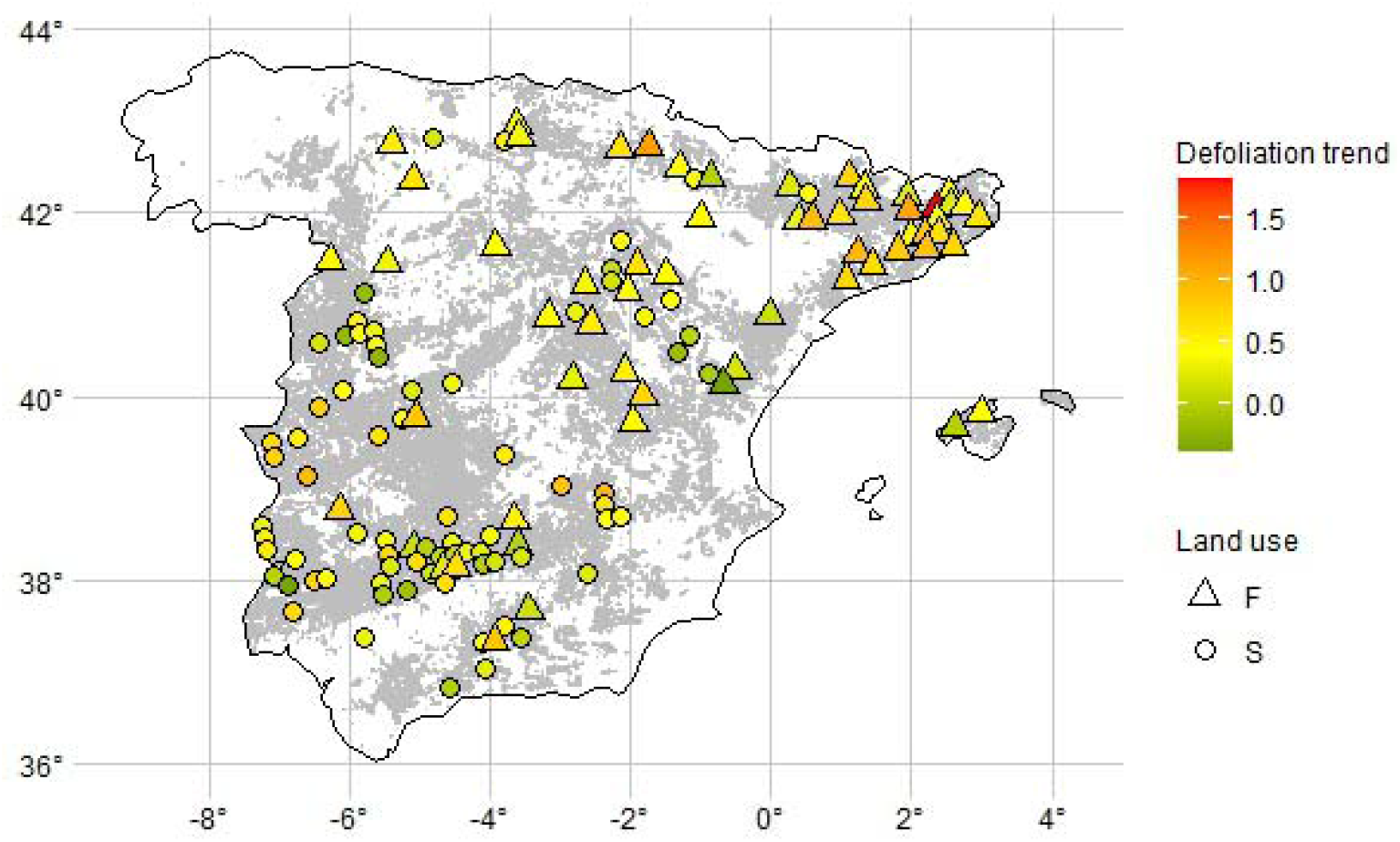
Crown tree defoliation trend over the 1987-2014 study period (% year-1 plot-1; in color) for the 134 savanna (circles) and forest (triangles) ICP holm oak plots across the distribution area of holm oak in Spain (grey area). The holm oak distribution map was provided by GIS-FOREST INIA (Ministerio de Medio Ambiente, y medio Rural y Marino, 2009).

**Figure 4.**
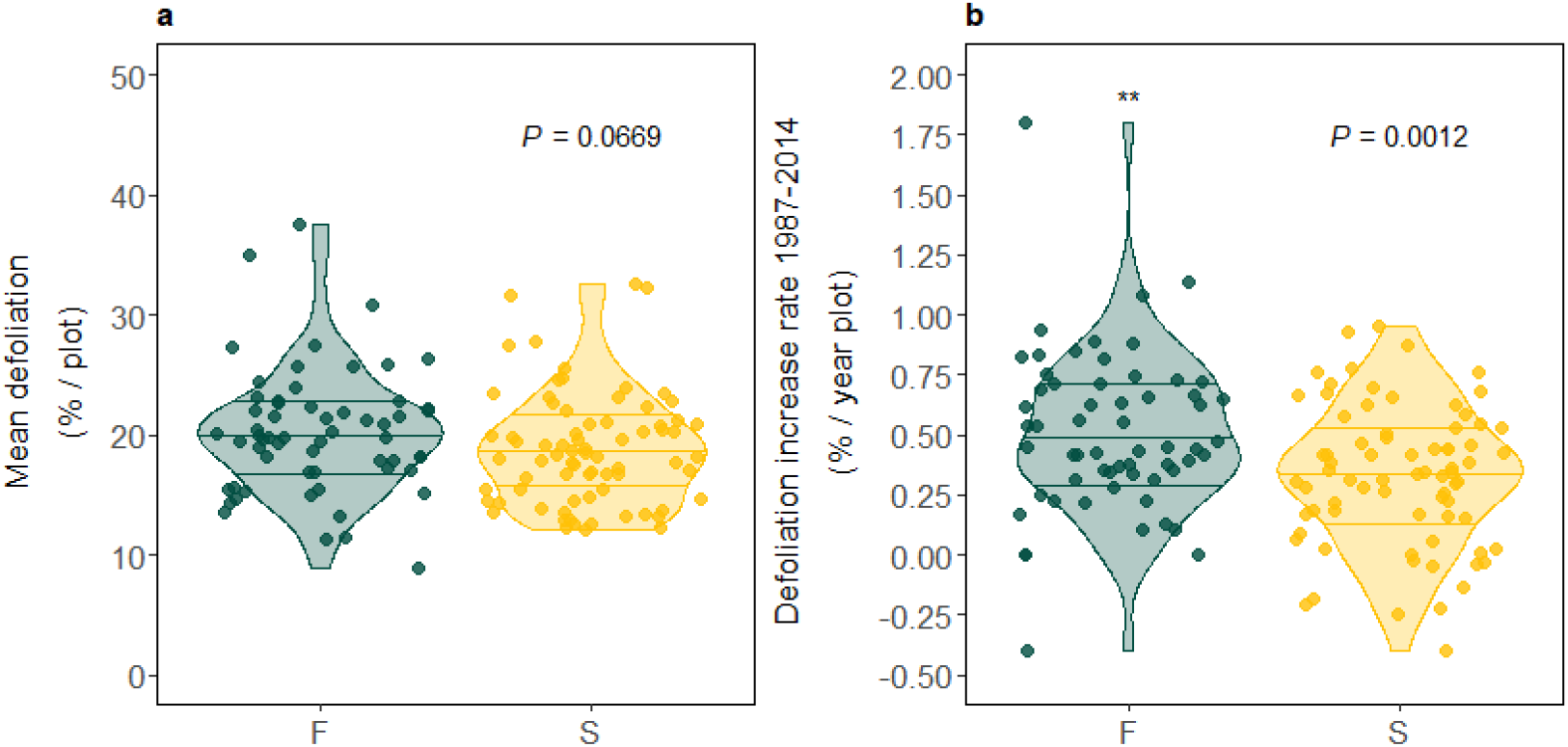
Violin plots of a) mean and b) trend of crown holm oak defoliation in savannas (S; yellow; n = 74) and forests (F; green; n = 60) over the 1987-2014 period. Asterisks represent significant differences (p-value < 0.05) between the two land-use types.

The results of the PCA with the topographic variables showed that the first two axes (i.e., principal components, PCs) explained 51.4% of the total variance (Fig. 5). The first axis (PC1) explained 29.7% of the total variance and was dominated by the slope (mean and variance), the altitude variance, and the mean of the topographic wetness index (twi.mean) (Fig. 6a), with mean slope and TWI being the highest negative and positive loadings, respectively (Fig. 6b). The second axis (PC2) explained 21.7% of the total variance and the variables that contributed the most were variance of the topographic wetness index (twi.var) and mean values of the planform (plancurv.mean) and profile (profcurv.mean) terrain curvatures (Fig. S1). Land-use types were differently distributed across the PC1 values. While forests (green ellipse in Fig. 5) covered the whole range of PC1 values, savannas were associated with more positive PC1 values. Overall, PCA results pointed out the link between land-use types and local topographic characteristics, the latter represented by PC1 with negative and positive values representing steeper and flatter land surfaces, respectively.

**Figure 5.**
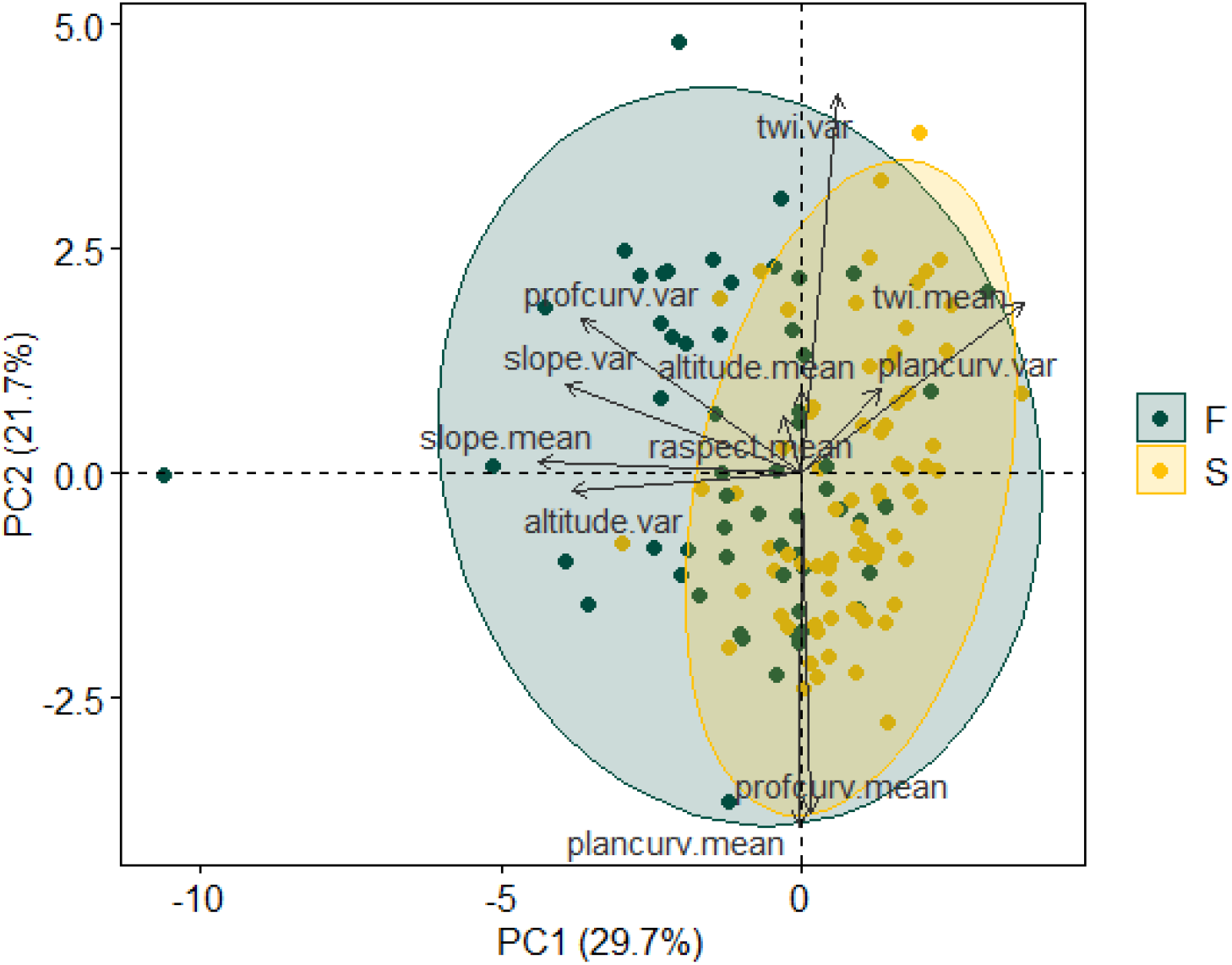
Biplot of the Principal Component Analysis (PCA) results for the topographic variables considered in this study, which correspond to mean (denoted as ‘.mean’) and variance (denoted as ‘.var’) values of altitude, slope, profile curvature (profcurv), planform curvature (plancurv), and topographic wetness index (twi), and reclassified mean aspect (raspect.mean). Dots represent the different ICP holm oak plots with colors denoting land-use types (S, savanna in yellow, and F, forest in green). Colored areas identify the general distribution of these land-use types over the first two principal components.

**Figure 6.**
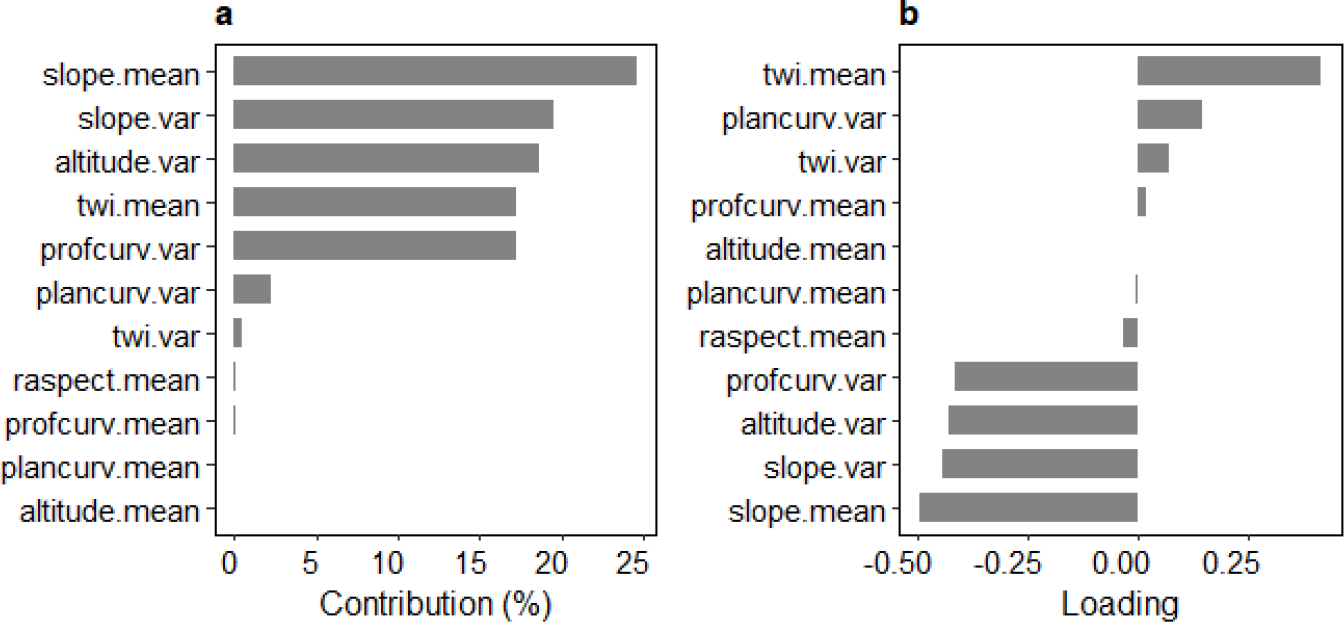
Bar plots of the first component’s (PC1) contributions (a) and loadings (b) of the twelve variables considered in the Principal Component Analysis (PCA).

The results of Spearman correlation analyses allowed the identification of the temporally aggregated seasonal climatic anomalies that correlated the most with defoliation trends (Fig. S2; ρ ≥ 0.20; p-value < 0.05) and did not show high correlation among them (Fig. S3; ρ ≤ 0.50; p-value < 0.05). These climatic anomalies were the median of precipitation anomaly over the previous winter, the mean of precipitation anomaly over the previous summer, the mean of the temperature anomaly over the previous spring, and the 28-years trends of minimum and maximum temperatures over the previous summer and autumn, respectively. Hence, aridity index and these climatic variables were included in the full model as well as their interactions with topography (PC1; Table S3). The selected defoliation model (Adj. R2 = 0.169, AIC = 62.84023, p-value = 0.00007465) obtained through backward and forward stepwise AIC fulfilled the residuals’ normality and heteroscedasticity assumptions. Coordinates were not included as explanatory variables as residuals did not show spatial autocorrelation (Moran’s I test for distance-based autocorrelation p-value = 0.3705). Among the predictors, only the interaction between topography (PC1) and the mean anomaly of precipitation over the previous summer showed a significant effect (p-value = 0.01; Table S3). However, other predictors that showed a marginal effect on defoliation trends (p-values = 0.05; Table S3) were the trend of the maximum temperature anomaly over the previous autumn and the interaction between topography (PC1) and the trend of the minimum temperature anomaly over the previous summer.

The model results suggest that the relationship between multidecadal trends of holm-oak defoliation and summer precipitation anomaly depends on local topography (Fig. 7). Those ICP plots located at steeper terrain with potentially lower water holding capacity (i.e., negative values of PC1) showed a negative relationship between defoliation trends and summer precipitation anomalies. Under these topographic conditions, a higher occurrence of summer droughts, depicted as negative values of the summer precipitation anomaly, was associated to higher defoliation trends. On the contrary, a positive relationship was only found for those plots with flatter topography (i.e., positive values of PC1), meaning that drier summers were related to lower defoliation trends.

**Figure 7.**
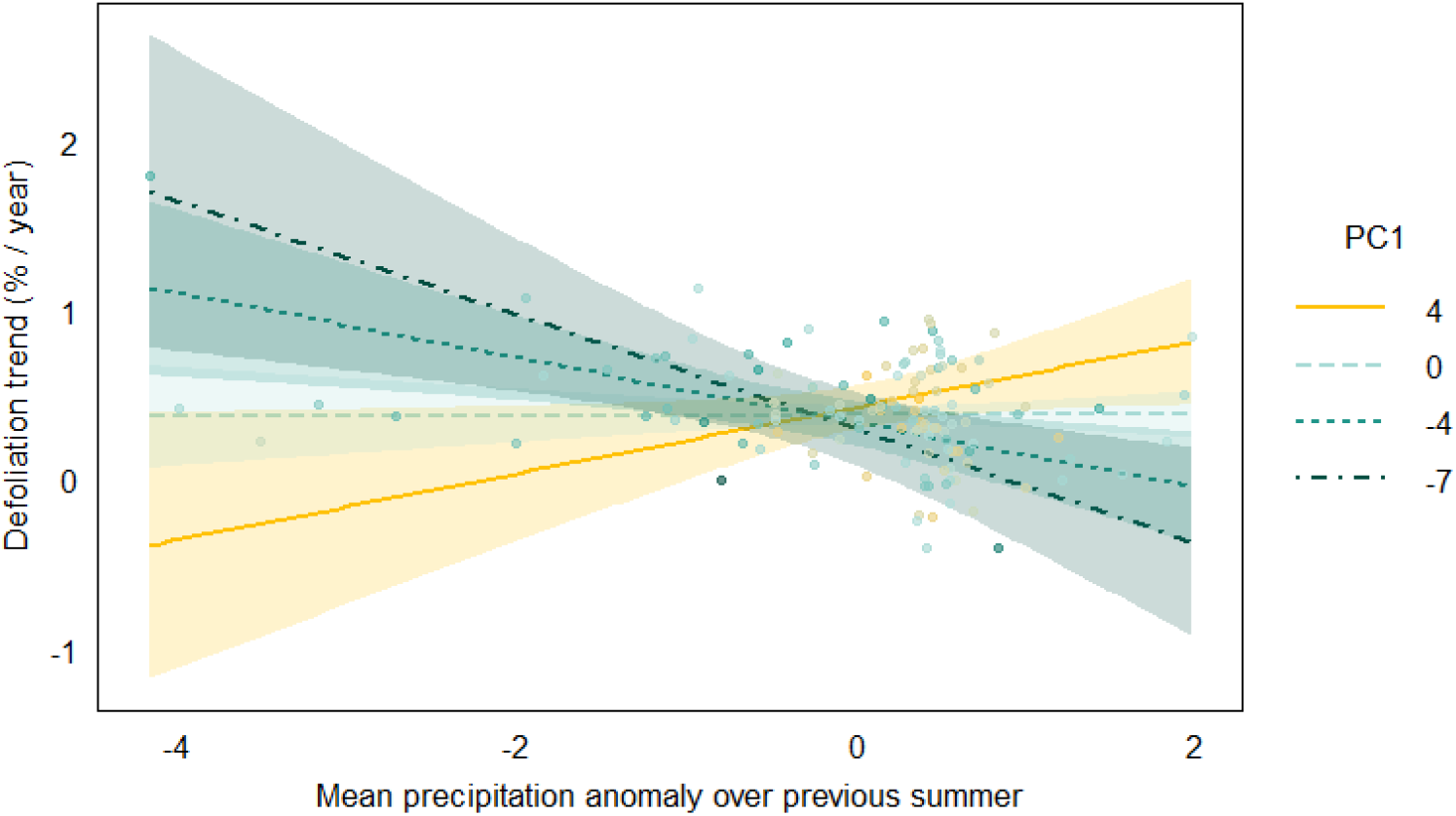
Multiple regression model results showing the multidecadal defoliation increase rates as a function of the interaction between the mean precipitation anomaly over the previous year with topography represented as PC1, with more positive values. Solid lines represent fitted data, while shaded areas correspond to the corresponding 95% confidence interval.

## 4. Discussion

Our work proposes an approach to perform multidecadal, regional, and observational assessments of forest decline that explicitly include topography as a modulator of tree climate sensitivity. Specifically, we investigated holm oak decline in Spain from 1987 to 2014 by using crown defoliation annual surveys performed under the European ICP Forests program. This data was collected at 134 plots where the principal land-use types (forest vs savanna) are represented, together with the topographic and climatic variability that define the area of distribution of this species within the Spanish territory. Our results show that holm oak defoliation has increased over the analyzed period, being this increase rate higher in forests compared to savannas. Moreover, we have detected that the interaction between topography and summer precipitation anomalies can partially explain trends of holm oak decline in the Mediterranean region. While a higher occurrence of dry summers increases defoliation trends in complex terrains where forests dominate, an inverse relationship was found at flatter terrains where savannas are located.

The jointly influence of climate extremes, land use, and topography has hardly been considered when trying to understand holm oak decline macro-dynamics by means of any of its proxies (i.e., tree growth, defoliation, or mortality). However, there are few studies that evaluate the role of some of these abiotic factors separately. Regarding the effect of land use, which is used here as a term that includes both forest structure and management, Gazol et al., (2020) noticed higher holm oak growth rates in savannas compared to forests distributed across Spain. They suggest that denser stands with smaller trees are subjected to higher water competition lowering their capacity to cope with droughts or heat waves. Their results support our finding of greater defoliation increase rates in forests compared to savannas (Fig. 4). Furthermore, water stress vulnerability in holm oak forest stands has been suggested to be exacerbated by land abandonment as it promotes natural regeneration through resprouting, successional vegetation growth and inter-specific water competition (Gentilesca et al., 2017; Martínez-Vilalta et al., 2012). This reinforces the growing evidence of the suitability of forest adaptive management as a way to improve the ecosystem health and resilience against climate extremes (Domingo et al., 2020; Dwyer et al., 2007; Galiano et al., 2012; Rodríguez-Calcerrada et al., 2011). Potential adaptive management strategies, such as thinning or extensive grazing, could be implemented particularly in high-density holm oak forests with abrupt topography and low terrain-driven soil moisture where traditional coppicing practices have been abandoned.

According to the PCA results, the land use types considered in our study are linked to specific local topographic characteristics (Fig. 5). On one hand, savannas are usually located at flatter terrains with deeper soils (Fig. 5,6) where roots can easily access water table levels (Moreno et al., 2005; Rolo & Moreno, 2012). On the other hand, only holm oak forests can be found at steeper areas with lower terrain-driven soil water availability (López et al., 2001). Generally, local topographic characteristics such as slope aspect and slope position are known to influence soil moisture availability in mountainous terrain (Stephenson, 1990). In this regard, previous research has demonstrated how topography can modulate vegetation responses to drought (Esteban et al., 2021; Hawthorne & Miniat, 2018; Paz-Kagan et al., 2017). These results obtained from local studies are in accordance with ours obtained regionally over three decades. Our model suggests that the most determinant factor influencing multidecadal defoliation trends was the interaction between topography and summer precipitation anomaly rather than climatic variables alone (Table S4). Concretely, a higher occurrence of drier summers was related to higher defoliation increase rates only for ICP plots located at steeper terrain conditions (i.e., negative PC1 values in Fig. 7), where abrupt topography (i.e., greater slopes and lower terrain-driven soil moisture) may limit soil water retention thus amplifying water stress and thermal exposure. This topography-climate interaction could mechanistically explain our finding of greater defoliation increase rates in forests compared to savannas (Fig. 4). These results are aligned with those by Galiano et al. (2012), who found a positive relationship between several decline indicators (i.e., canopy browning, and tree and stem mortality) and altitude in a holm oak forest located in NE Spain and suggested that, in this case, the negative effect of wind and radiation exposure on the ridge tops prevailed over the effect of cooler temperatures at higher altitudes.

Conversely, previous research primarily focused on holm oak savannas indicates no clear effect of micro-topographical features on defoliation at neither local (Corcobado et al., 2013) or regional scales (Sánchez-Cuesta et al., 2021). Regarding climate, Sánchez-Cuesta et al. (2021) identified aggregated summer SPEI over the prior 18 months and mean annual temperature as the two best predictors of holm oak annual defoliation in savannas distributed across southwestern Spain, showing negative and positive relationships, respectively. In contrast, we found that a higher number of drier summers were related with lower defoliation increase rates in ICP plots situated at flatter terrains with potentially higher topography-driven soil moisture conditions (i.e, positive values of PC1; Fig. 7), where most of the analyzed savannas are located (Fig. 5). Based on previous research, we suggest that this unexpected result might be explained by the fact that Phytophtora cinamonii (hereinafter Phytophtora spp.), the primary holm oak pathogen, needs soil saturation conditions to proliferate (Gisi et al., 1980; Sterne et al., 1977). Accordingly, we found a positive relationship between tree health and dryness only for those locations more susceptible of waterlogging (i.e., higher PC1 values associated to higher TWI values; Fig 6). Indeed, a recent study based on a greenhouse experiment have shown that water stress can be beneficial for Quercus suber seedlings as it can reduce the Phytophtora spp. root damage (Homet et al., 2019). However, although the ICP database provide information about the cause of damage from 2005, none of the analyzed ICP plots here has been linked to Phytophtora spp. infection, probably due to the great difficulty to identify it.

In order to better understand spatiotemporal patterns of holm oak decline, future research may include spatial explicit soil characterization data (e.g., SoilGrids) with greater spatial resolution, particularly on soil texture given its link with Phytophtora spp. proliferation (Corcobado et al., 2013). Similarly, spatial explicit quantitative information on forest structure would enhance the predictive power of our model. Here, the time series of European tree canopy cover initiated in 2012, first with a spatial resolution of 20 m and further on of 10 m will become a powerful data source to include tree cover in future analysis. Furthermore, although our study has a relatively large scale, it does not cover the whole distribution area of Quercus ilex, which spans from Portugal and Morocco to the Aegean Islands and western Turkey expanding northward up to northern Italy and France (de Rigo and Caudullo, 2016). International georeferenced assessments are challenging as land use, climate and topographical maps at sufficient spatial and temporal resolutions may be in different formats, repositories or even not be publicly accessible. Nevertheless, our results contribute to the current understanding of Mediterranean forest decline as it probes, based on the categorization of decline factors developed by Manion (1981), the great relevance of topography together with land use as very influential predisposing factors that can modulate both how climate extremes (as inciting factors) affect tree health, as well as the link between these inciting factors with other contributing factors, such as pathogen infection (Gentilesca et al., 2017). Our results are in accordance with recent research that acknowledges the outstanding role of local topography in explaining the spatial variability of forest resilience (Bramer et al., 2018; Carnicer et al., 2021; Kopecký et al., 2021; Kopecký & Čížková, 2010; Petroselli et al., 2013), as it regulates tree water availability together with thermal, radiation and wind exposure. Overall, local stand and topographic characteristics shape forest microclimate that ultimately drives the tree response to warmer and drier climatic conditions (Zellweger et al., 2020).

## 5. Conclusions

Contrary to previous research usually constrained to a certain region, land use type or decline factor, our study provides a comprehensive conceptual framework to: (1) investigate holm oak decline in the widest possible spectrum of uses, topography and climate that define the distribution of this species of enormous ecological importance in the Mediterranean basin; and (2) understand, using unprecedented spatial (countrywide) and temporal (multi-decadal) scales, the sensitivity of Iberian holm oaks to key abiotic factors and their interactions in space and time. Our results contribute to the incipient but growing scientific trend that acknowledges the influence of local topography on forest functioning, as it can shape forest vulnerability against climate extremes. The present work offers guidance to identify potential holm oak decline hotspots and design proper spatial-explicit measures to improve the health and future stability of Mediterranean oak ecosystems in current and future climate change scenarios. Broadly speaking, our study shows how the future management of holm oak stands will be key to guaranteeing their survival, especially in the topographically more complex areas where the effects of climate change can be much more damaging.

## Acknowledgments

This work was supported by a Juan de la Cierva-Incorporación postdoctoral contract IJC2020-045630-I and the MANAGE4FUTURE project (TED2021-129499A-I00) both funded by MCIN/AEI /10.13039/501100011033 and the European Union NextGenerationEU/PRTR. Ana-Maria Hereş was financed by the REASONING (PN-III-P1-1.1-TE-2019-1099) project through UEFISCDI (link; Romanian Ministry of Education and Research). This research was supported by the BERC 2018-2021 (Basque Government), and BC3 María de Maeztu Excellence Accreditation 2018-2022, Ref.MDM-2017-0714 (Spanish Ministry of Science, Innovation and Universities).

## Data availability statements

Databases utilized in this study are all publicly available at the sources in indicated in material and methods and supplementary materials sections.

