## Supplementary materials for "Topography modulates climate sensitivity of multidecadal trends of holm oak decline"

**Table S1.** Main characteristics of the geospatial and survey datasets used in this study.

| Cartographic & RS Databases | Variables | Spatial resolution | Temporal resolution | Start date | End  date |
| --- | --- | --- | --- | --- | --- |
| ICP forests survey | Tree defoliation (%) | In situ observations structured on a 16x16 km grid | Annual | 1987 | 2014 |
| SIOSE | Land use  (% cover type) | 1:25000 | 2005, 2011, 2014 | 2005 | 2014 |
| Digital Elevation Model (MDT05) | Elevation | 5 m | - | - | - |
| Global Aridity Index (AI) Database v2 | AI^1^ | 1km |  |  |  |
| Spain02v05 | Precipitation, mean, max and min temperature | 10 km (0.1 deg) | Monthly | 01/01/1950 | 31/12/2015 |

^1^Mean AI values for the 1970-2000 period.

**Table S2.** Filters applied to each raster file associated to each topographic variable used in this study to assure plausible ranges prior to the extraction of zonal statistics.

| Variable | Range | Units |
| --- | --- | --- |
| Altitude | ≥ 0 | meters |
| Slope | [0, 90] | degrees |
| Aspect | [0, 360] | degrees |
| Profile curvature | (-15, 15) | dimensionless |
| Planform curvature | (-15, 15) | dimensionless |
| Topographical Wetness Index | ≥ 0 | dimensionless |

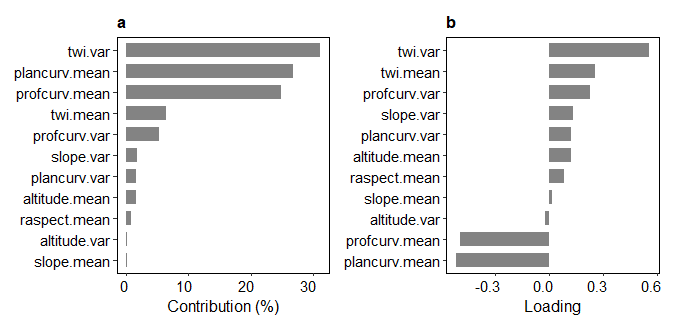

**Figure S1.** Bar plots of the contributions (**a**) and loadings (**b**) of the twelve variables considered in the Principal Component Analysis (PCA) for the second principal component (PC2).

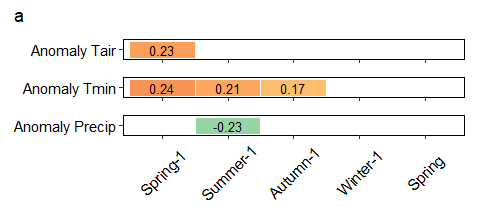

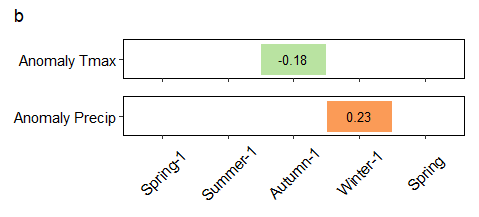

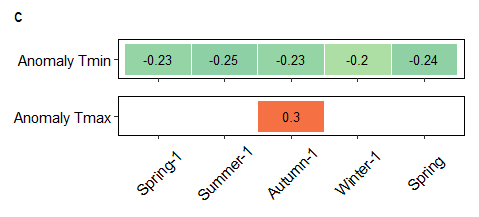

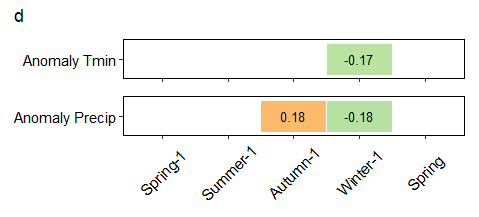

**Figure S2.** Significant Spearman correlation coefficients (p-value < 0.05) between defoliation trends and (**a**) mean, (**b**) median, (**c**) trend, and (**d**) skewness of seasonal anomalies of mean, minimum and maximum temperature and precipitation from spring of the previous-to-defoliation measurements years (-1) to spring of the current-to-defoliation measurements years for the 134 ICP holm oak plots analyzed in this study. The four panels correspond to correlation matrices performed with the mean, median, Sen´s slope and skewness values of climatic seasonal anomalies over the study period (1987-2014). Colors depict the magnitude and sign of the correlation while empty cells representing non-significant relationships.

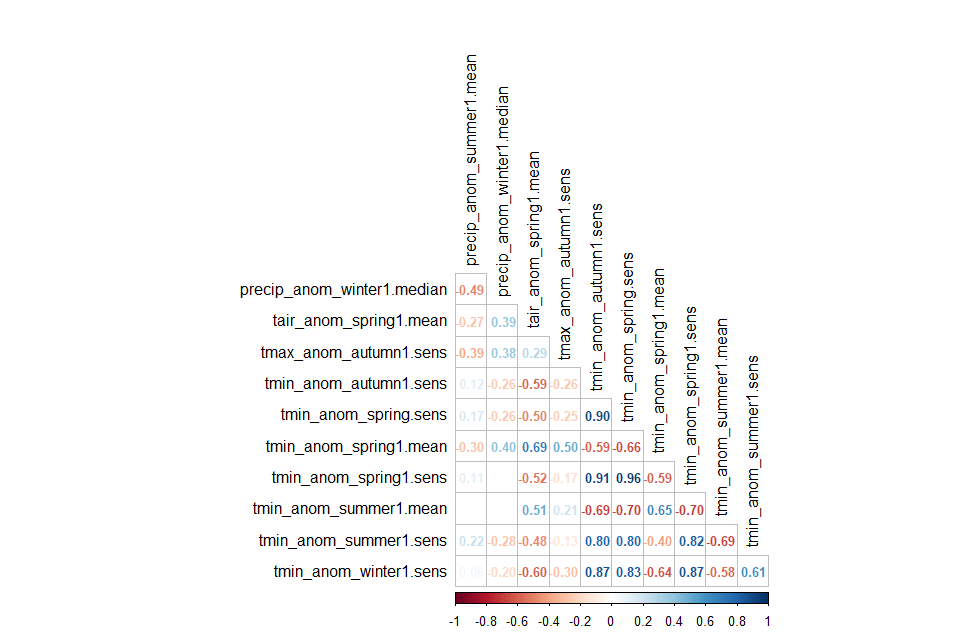

**Figure S3.** Significant Spearman correlation coefficients (p-value < 0.05) among temporally aggregated seasonal anomalies highly correlated with defoliation trends. Colors depict the magnitude and sign of the correlation while empty cells represent non-significant relationships.

**Table S3.** Model names and formulations.

| Model name | Formulation |
| --- | --- |
| Full model | defol.trend =  precip_winter1_median*pc1 +  precip_summer1_mean*pc1 +  tair_spring1_mean*pc1 +  tmin_summer1_trend*pc1 +  tmax_autumn1_trend*pc1 +  precip_summer1_mean*tmin_summer1_trend +  ai |
| Selected model | defol.trend =  precip_winter1_median +  pc1 +  precip_summer1_mean +  tmin_summer1_trend +  tmax_autumn1_trend +  pc1:precip_summer1_mean +  pc1:tmin_summer1_trend |

**Table S4.** Coefficient estimates together with associated confidence intervals (CI; at 95% significance level) and p-values for the selected model.

| Predictors | Estimates | CI | p |
| --- | --- | --- | --- |
| (Intercept) | 0.38 | 0.33 – 0.44 | **<0.001** |
| precip winter1 median | 0.05 | -0.01 – 0.11 | 0.081 |
| pc1 | 0.01 | -0.02 – 0.04 | 0.354 |
| precip summer1 mean | 0.00 | -0.07 – 0.07 | 0.953 |
| tmin summer1 trend | -0.04 | -0.09 – 0.02 | 0.211 |
| tmax autumn1 trend | 0.06 | -0.00 – 0.12 | 0.052 |
| pc1 × precip summer1 mean | 0.05 | 0.01 – 0.08 | **0.009** |
| pc1 × tmin summer1 trend | 0.03 | -0.00 – 0.06 | 0.053 |
